# Inhibition of mammalian mtDNA transcription paradoxically activates liver fatty acid oxidation to reverse diet-induced hepatosteatosis and obesity

**DOI:** 10.1101/2023.09.22.558955

**Authors:** Shan Jiang, Taolin Yuan, Laura S Kremer, Florian A Rosenberger, Fynn M Hansen, Melissa Borg, Diana Rubalcava-Gracia, Mara Mennuni, Roberta Filograna, David Alsina, Jelena Misic, Camilla Koolmeister, Lipeng Ren, Olov Andersson, Anke Unger, Tim Bergbrede, Raffaella Di Lucrezia, Rolf Wibom, Juleen R Zierath, Anna Krook, Patrick Giavalisco, Matthias Mann, Nils-Göran Larsson

**Affiliations:** Department of Medical Biochemistry and Biophysics, Karolinska Institutet, Stockholm, Sweden; Department of Proteomics and Signal Transduction, Max-Planck Institute of Biochemistry, Martinsried, Germany; Department of Physiology and Pharmacology, Section for Integrative Physiology, Karolinska Institutet, Stockholm, Sweden; Department of Cell and Molecular Biology, Karolinska Institutet, Stockholm, Sweden; Lead Discovery Center GmbH, Otto-Hahn-Str. 15, Dortmund, Germany; Centre for Inherited Metabolic Diseases, Karolinska University Hospital, Stockholm, Sweden; Department of Molecular Medicine and Surgery, Section for Integrative Physiology, Karolinska Institutet; Metabolomics Core Facility, Max Planck Institute for Biology of Ageing, Cologne, Germany

## Abstract

The oxidative phosphorylation (OXPHOS) system in mammalian mitochondria plays a key role in harvesting energy from ingested nutrients^1, 2^. Mitochondrial metabolism is very dynamic and can be reprogrammed to support both catabolic and anabolic reactions, depending on physiological demands or disease states^3, 4^. Rewiring of mitochondrial metabolism is intricately linked to metabolic diseases^5, 6^ and is also necessary to promote tumour growth^7–11^. Here, we demonstrate that *per oral* treatment with an inhibitor of mitochondrial transcription (IMT)^11^ shifts whole animal metabolism towards fatty acid oxidation, which, in turn, leads to rapid normalization of body weight, reversal of hepatosteatosis and restoration of glucose tolerance in mice on high-fat diet. Paradoxically, the IMT treatment causes a severe reduction of OXPHOS capacity concomitant with a marked upregulation of fatty acid oxidation in the liver, as determined by proteomics and non-targeted metabolomics analyses. The IMT treatment leads to a marked reduction of complex I, the main dehydrogenase that feeds electrons into the ubiquinone (Q) pool, whereas the levels of electron transfer flavoprotein dehydrogenase (ETF-DH) and other dehydrogenases connected to the Q pool are increased. This rewiring of metabolism caused by reduced mtDNA expression in the liver provides a novel principle for drug treatment of obesity and obesity-related pathology.

## Main

The first attempts to target mitochondria to treat obesity were reported in the 1930s when more than 100,000 individuals were treated with the uncoupler dinitrophenol (DNP)^12–14^. Although DNP treatment increased the metabolic rate and reduced obesity, serious side-effects prevented it from becoming an established treatment^14, 15^ and research is ongoing to find new uncouplers with better properties for treatment of obesity^16^ and cancer^17^. Metformin provides an alternate way to inhibit OXPHOS and this mild complex I inhibitor is widely used as an antidiabetic medication^18–20^ and also protects against cancer^21–24^. The possible connection between beneficial metabolic effects and anti-cancer activity of drugs targeting mitochondria prompted us to investigate whether IMT treatment, which is known to impair tumor metabolism and growth in mouse models^11^, also may have beneficial metabolic effects. Treatment of tumor cell lines with IMT induces a dose-dependent impairment of OXPHOS and cellular metabolic starvation with progressively reduced levels of a range of critical metabolites and eventually cell death^11^. Despite the drastic effects on metabolism in cancer cell lines and cancer xenografts, treatment of whole animals is well tolerated^11^. We therefore decided to test the hypothesis that IMT-treatment to impair the OXPHOS capacity in whole animals may induce beneficial metabolic effects in normal and metabolically challenged mice.

### IMT treatment protects against HFD-induced obesity

To evaluate the metabolic effects of inhibited mtDNA transcription, male C57BL/6N mice at the age of four weeks were randomly chosen to be fed standard chow diet or HFD for eight weeks. Thereafter, the two groups were subdivided for *per oral* treatment (gavage) with either IMT (LDC4857, 30 mg/kg) or vehicle for four weeks while continuing the respective diets (Fig. 1a). The IMT compound used in this study was developed within an optimization program based on the structurally closely related IMT1B compound, which previously has been used for *per oral* treatment (100 mg/kg) of cancer xenografts^11^. IMT treatment of mice on HFD causes a rapid marked reduction of body weight after one week, with a cumulative weight loss of ∼7 g after four weeks (Fig. 1b). Measurements of body composition with non-invasive magnet resonance imaging (EchoMRI-100^TM^) after four weeks of IMT treatment showed markedly reduced fat mass without any change of lean mass (Fig. 1c). We performed hematoxylin-eosin (H&E) staining of tissue sections of epididymal white adipose tissue (eWAT) and found that HFD leads to the expected formation of large, lipid-filled adipocytes (Extended Data Fig. 1a). In contrast, IMT treatment during HFD led to a drastic decrease in adipocyte size (Extended Data Fig. 1a), consistent with the observed decrease of body weight (Fig. 1b) and whole body fat content (Fig. 1c).

**Fig. 1:**
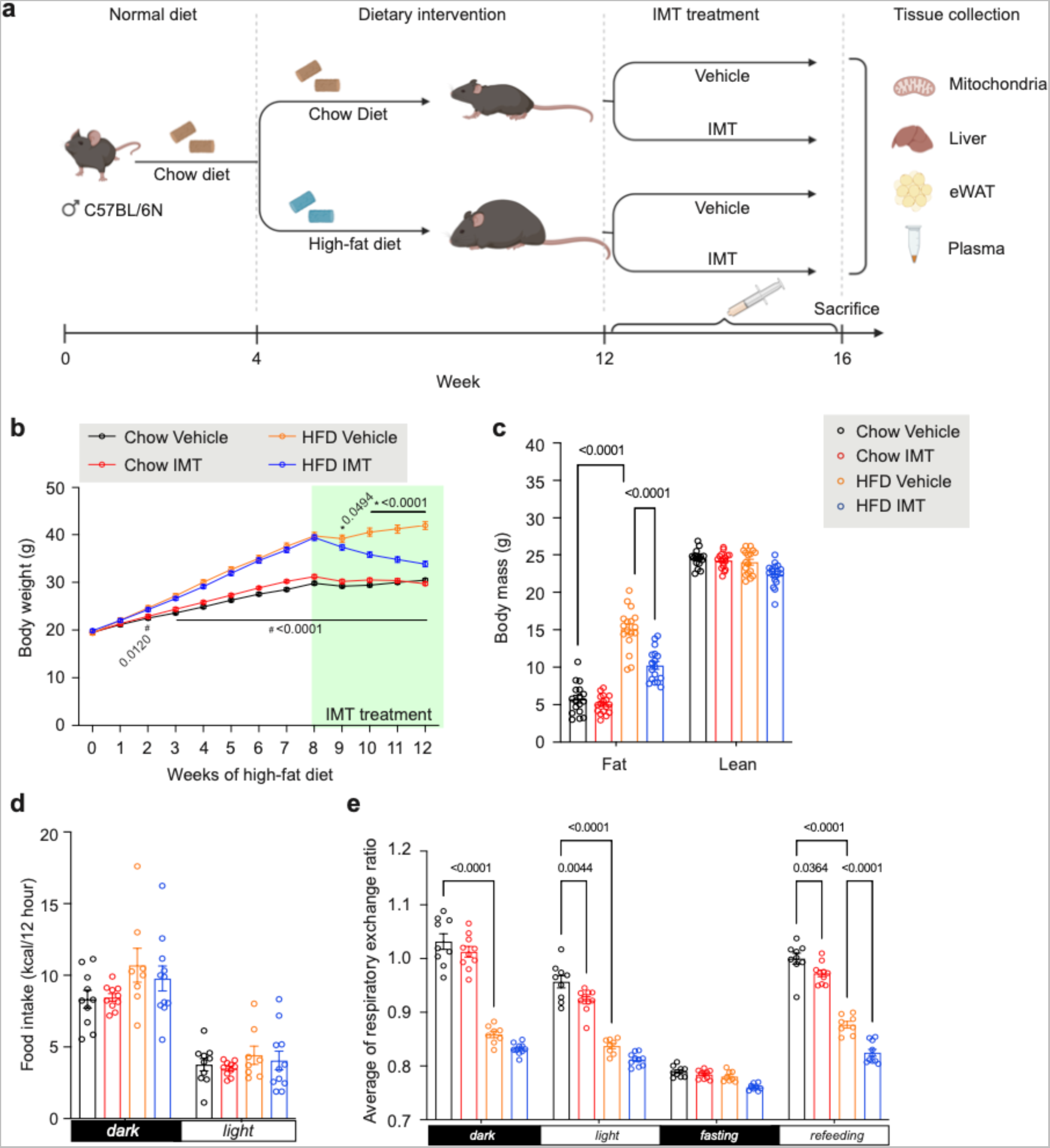
IMT treatment prevents diet-induced obesity and improves glucose homeostasis. **a**, Experimental strategy for diet intervention and IMT treatment. Male C57BL/6N mice at the age of four weeks were randomly fed either chow diet or HFD for eight weeks. Thereafter, the diet was continued, and mice were *per orally* treated with IMT (30 mg/kg) or vehicle for four weeks. Six independent cohorts of mice were used in this study, total mice n = 260. **b,** Body weight in mice on chow diet or HFD treated with vehicle or the IMT compound. n = 22 mice per group. Data are presented as mean ± SEM. Statistical significance was assessed by a two-way ANOVA with Tukey’s test for multiple comparisons. ∗ indicates a significant difference between HFD IMT and HFD Vehicle; ^#^ indicates a significant difference between Chow Vehicle and HFD Vehicle. P-values are shown in the figure. **c**, Body composition showing fat mass and lean mass after four weeks of IMT treatment. n = 17 mice per group. **d, e,** Measurement of whole-body metabolism during the fourth week of gavage treatment with vehicle or IMT compound by using the Oxymax/Comprehensive Lab Animal Monitoring System (CLAMS). Food intake during the fourth day (**d**). The average of respiratory exchange ratio (RER) over 42 hours during the light and dark cycles (**e**). Chow vehicle n = 10, Chow IMT n = 10, HFD vehicle n = 8, and HFD IMT n = 11 mice. For **c**-**e**, data are presented as mean ± SEM. Statistical significance was assessed by a two-way ANOVA with Tukey’s test for multiple comparisons. P-values are shown in the figure.

We next assessed whole body energy homeostasis in mice on chow diet or HFD treated with vehicle or the IMT compound by using the Oxymax/Comprehensive Lab Animal Monitoring System (CLAMS). The four groups of mice were subjected to five continuous days of CLAMS analysis during the fourth week of gavage treatment with vehicle or IMT compound. The first three days were used to acclimate the animals to the CLAMS system, followed by measurements during the fourth day. Day five included a 12-hour period of fasting followed by six-hour of refeeding. Importantly, IMT treatment did not alter food intake (Fig. 1d) or physical activity (Extended Data Fig. 1b). However, the IMT-treated mice on HFD showed enhanced oxygen consumption during both the light and dark cycle (Extended Data Fig. 1c), consistent with increased mitochondrial metabolism. During HFD feeding, the respiratory exchange ratio (RER) was decreased to ∼0.8 (Fig. 1e, Extended Data Fig. 1d), signifying a preference for fat as a metabolic fuel substrate, whereas mice on the standard chow diet had a RER of ∼0.9 – 1.1. Upon refeeding after fasting, the IMT treatment resulted in a lower RER in comparison with vehicle treatment, regardless of the diet (Fig. 1e, Extended Data Fig. 1d), pointing to drug-induced activation of fat metabolism. These data provide evidence that IMT treatment reverses HFD-induced obesity by promoting the use of fat as an energy source at the organismal level.

### IMT treatment reverses diet-induced glucose intolerance

Obesity is a main risk factor for type 2 diabetes, and we therefore assessed glucose homeostasis in mice on HFD. We found normal fasting blood glucose levels accompanied by markedly increased fasting serum insulin levels (Fig. 2a, b). Furthermore, mice on HFD had pathological intraperitoneal glucose tolerance tests (ipGTT; Fig. 2c, d) with an increased peak concentration of serum insulin (Fig. 2e), consistent with a pre-diabetic state and insulin resistance. Glucose homeostasis was markedly improved when mice on HFD were treated with IMT compound for four weeks, i.e., the fasting blood glucose was reduced (Fig. 2a), the serum insulin levels were decreased (Fig. 2b) and the ipGTT responses were normalized Fig. 2c-e).

**Fig. 2:**
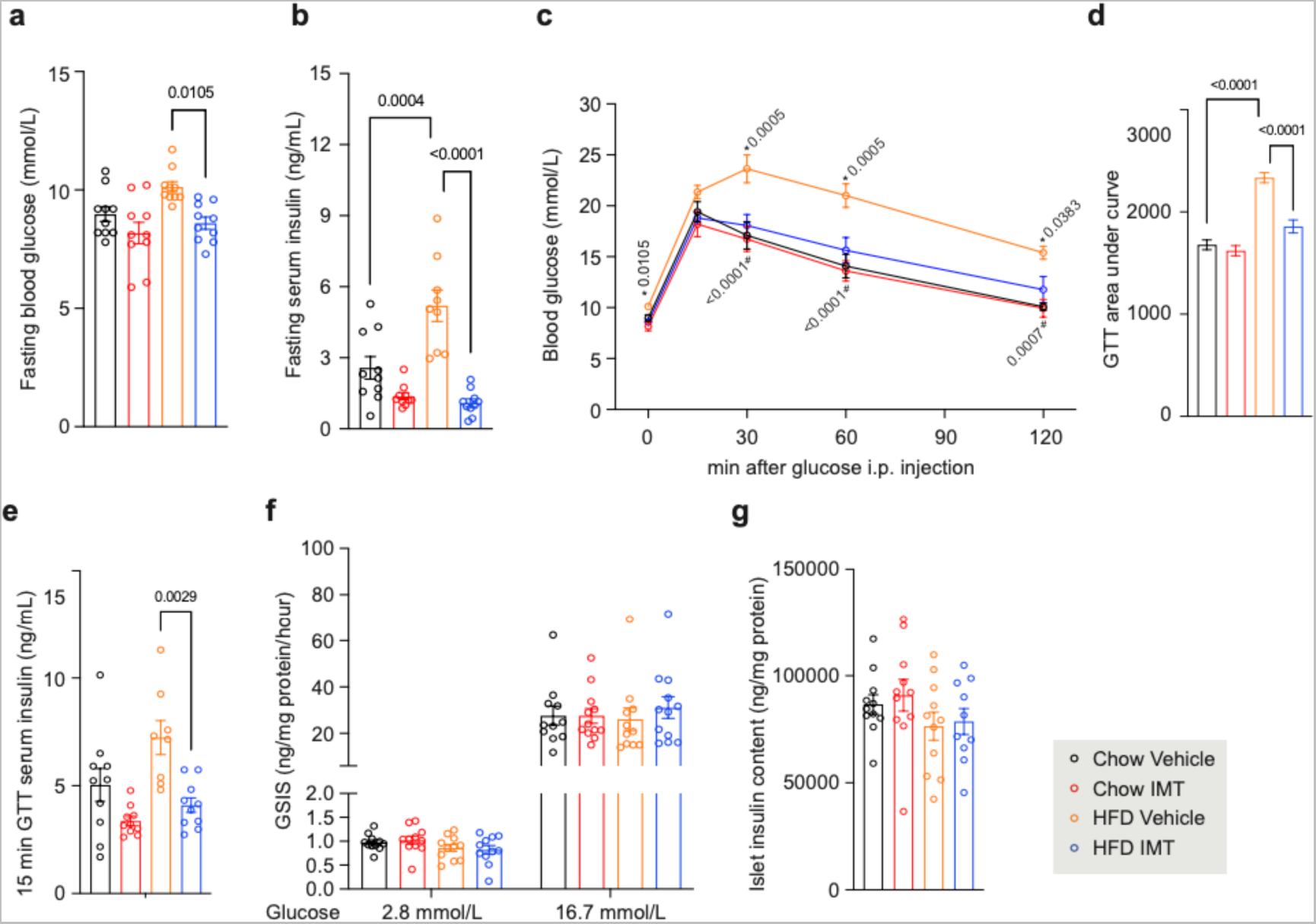
IMT improves glucose homeostasis and does not impair islet insulin secretion. **a**, **b**, Fasting blood glucose (**a**) and fasting serum insulin (**b**) levels in mice after four weeks of IMT treatment. n = 10 mice per group. **c, d**, Blood glucose levels (**c**) and the area under the curve (AUC, **d**) during intraperitoneal glucose tolerance tests (ipGTT) with 2 g/kg glucose in mice after four weeks of vehicle or IMT treatment. n = 10 mice per group. Data are presented as mean ± SEM. Statistical significance was assessed by a two-way ANOVA with Tukey’s test for multiple comparisons. ∗ indicates a significant difference between HFD IMT and HFD Vehicle; ^#^ indicates a significant difference between Chow Vehicle and HFD Vehicle. P-values are shown in the figure. **e,** Serum insulin levels at the 15-min of ipGTT. n = 10 mice per group. **f**, Ex vivo glucose-stimulated insulin secretion assays performed on isolated pancreatic islets. Glucose (2.8 mM and 16.7 mM) was added to the medium to recapitulate basal and glucose-stimulated insulin secretion conditions. Three independent GSIS assays were performed with three replicates per group. **g**, Islet insulin content. Three independent GSIS assays were performed with three replicates per group. All data are presented as mean ± SEM. Statistical significance was assessed by a two-way ANOVA with Tukey’s test for multiple comparisons. P-values are shown in the figure.

Transcription of mtDNA and OXPHOS function are essential for the stimulus-secretion coupling in pancreatic β-cells^25^. IMT treatment leads to reduced circulating insulin levels (Fig. 2b, e) and these findings prompted us to investigate whether insulin secretion in pancreatic β-cells is impaired. We performed *ex vivo* glucose-stimulated insulin secretion assays (GSIS) on isolated pancreatic islets and found that four weeks of IMT treatment did not impair insulin secretion or insulin biosynthesis regardless of the diet (Fig. 2f, g). The reduced circulating insulin levels and normalized glucose homeostasis in IMT-treated mice on HFD are thus likely explained by improved insulin sensitivity.

### Reversal of hepatic steatosis by IMT treatment in mice on HFD

We observed a massive macrovesicular steatosis in liver of mice on HFD (Fig. 3a). Remarkably, IMT treatment drastically reduced hepatosteatosis (Fig. 3a), leading to decreased lipid content (Fig. 3b) and reduced liver weight (Fig. 3c). We performed untargeted lipidomics to measure the lipid species in mouse liver (Fig. 3d). The HFD led to a massive accumulation of diglycerides and triglycerides in the liver, which was reversed after four weeks of IMT treatment (Fig. 3d). In contrast, the phospholipid and sphingolipid levels in liver were mainly affected by the diet and not markedly impacted by IMT treatment (Extended Data Fig. 2). The reduction of diglycerides and triglycerides in liver of mice on HFD treated with IMT was accompanied by improvement of liver function as measured by decreased aminotransferase levels in serum (Fig. 3e, f). The serum albumin levels were normal in all investigated groups (Fig. 3g). Taken together these data show that IMT-treatment can reverse diet-induced hepatosteatosis and normalize liver function.

**Fig. 3:**
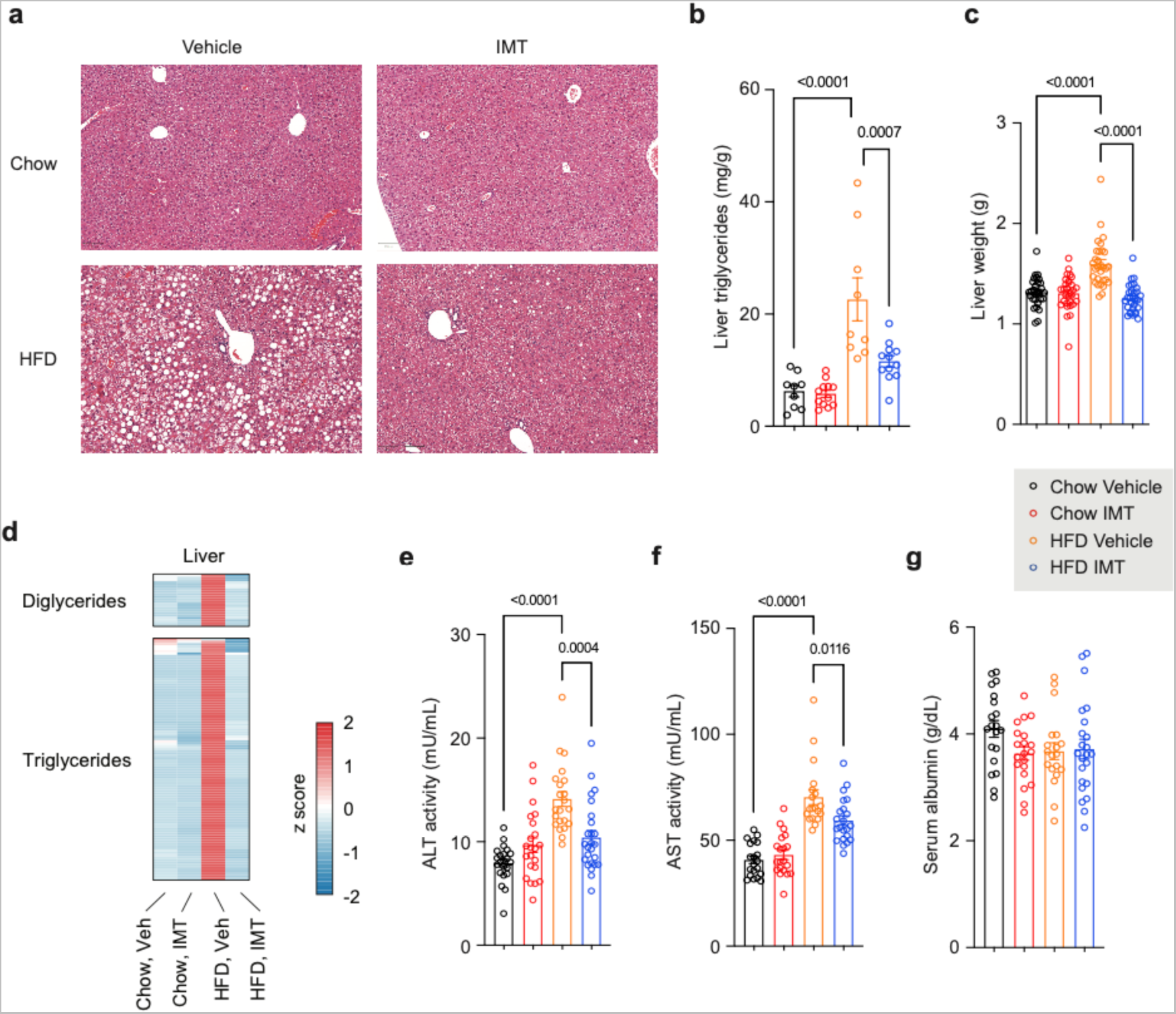
IMT treatment reverses hepatosteatosis. **a**, Representative images of H&E staining showing liver structure and morphology in mice on chow diet or HFD treated with either vehicle or IMT compound. Scale bar, 100 μm. n = 5 mice per group. **b**, Quantitative measurement of triglycerides in mouse liver after four weeks of IMT treatment. n = 12 mice per group. **c**, Liver weight in mice treated with vehicle or IMT for four weeks. n = 30 mice per group. **d**, The levels of diglycerides and triglycerides in mouse liver after four weeks of IMT treatment. Chow vehicle, Chow IMT and HFD vehicle n = 8 mice per group, HFD IMT n = 7 mice. **e-g**, The serum levels of alanine aminotransferase, ALT (**e**), aspartate aminotransferase, AST (**f**), and albumin (**g**) measured in mice after four weeks of vehicle or IMT treatment. n = 18 mice per group. For **b**, **c**, **e**-**g**, data are presented as mean ± SEM. Statistical significance was assessed by a two-way ANOVA with Tukey’s test for multiple comparisons. P-values are shown in the figure.

### IMT treatment preferentially inhibits mtDNA transcription in liver

IMT treatment resulted in marked reduction in levels of mtDNA-encoded transcripts (Fig. 4a) and mtDNA (Fig. 4b) in liver of mice on chow diet or HFD. The decrease of mtDNA is likely due to decreased formation of RNA primers needed for initiation of mtDNA replication, because IMT inhibits POLRMT which serves as the primase for mammalian mtDNA replication^26^. Treatment with IMT also resulted in moderate decrease of mtDNA-encoded transcripts and mtDNA levels in eWAT (Extended Data Fig. 3a, b), whereas there were no significant changes in skeletal muscle (Extended Data Fig. 3c, d). IMT treatment caused no reduction of mtDNA-encoded transcripts in heart and brown adipose tissue (BAT) (Extended Data Fig. 3e, f). To gain further insights into the differences in inhibition of mtDNA transcription between tissues, we measured IMT concentrations 24 hours after the last dose in mice treated with IMT for four weeks (Fig. 4c). The IMT concentration was much higher in plasma and liver than in heart, skeletal muscle, eWAT and BAT, which argues that the IMT compound is preferentially accumulated in liver due to the *per oral* route of administration and the first-passage effect. The skewed tissue distribution of IMT thus likely explains the marked liver-specific inhibitory effect on mtDNA transcription.

**Fig. 4:**
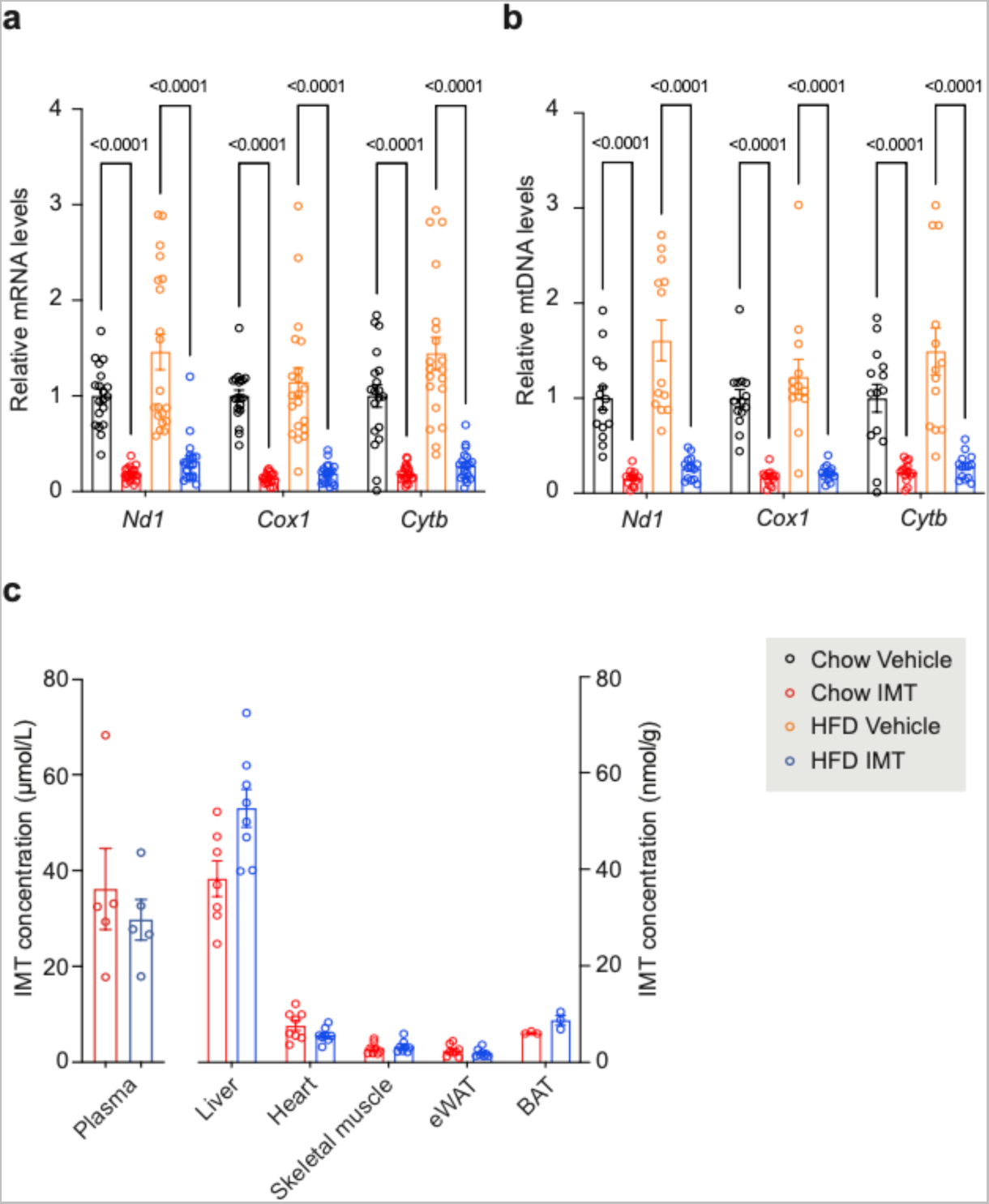
IMT treatment impairs mtDNA transcription. **a**, Mitochondrial transcript levels in liver after four weeks of IMT treatment. n = 20 mice per group. **b**, Levels of mtDNA in liver after four weeks of IMT treatment. n = 14 mice per group. **c**, IMT concentration in plasma and mouse tissues. Plasma n = 5 mice per group; liver Chow IMT n = 7 mice, HFD IMT n = 8 mice; heart, skeletal muscle, eWAT n = 8 mice per group; BAT n = 3 mice per group. For **a**-**c**, data are presented as mean ± SEM. Statistical significance was assessed by a two-way ANOVA with Tukey’s test for multiple comparisons. P-values are shown in the figure.

### Rewiring of OXPHOS pathways in IMT-treated liver

We used label-free quantitative proteomics to identify differentially expressed proteins in homogenates from liver tissue or ultra-purified liver mitochondria. In total, 4,408 proteins were identified in the liver tissue proteome of mice on chow diet or HFD, and IMT treatment caused a significant change in the levels of 15% - 20% of these proteins (FDR < 0.05, Extended Data Fig. 4a). A high proportion (68.7% at FDR < 0.05) of the proteins whose levels changed significantly after IMT treatment were classified as mitochondrial proteins according to MitoCarta 3.0^27^. We performed principal component analyses and found that changes in the total and mitochondrial liver proteome both were mainly determined by the IMT treatment (Extended Data Fig. 4b).

A detailed inspection of the OXPHOS subunits in the mitochondrial proteome showed that IMT treatment significantly decreased the levels of subunits of complex I, III, IV and the membrane portion (F_0_) of complex V, whereas subunits of the matrix portion (F_1_) of complex V were less affected or were increased (Extended Data Fig. 4c, Extended Data Fig. 5a). In contrast, the levels of subunits of complex II (succinate dehydrogenase) were increased (Extended Data Fig. 5a), consistent with the lack of mtDNA-encoded subunits in this complex. Western blot analyses confirmed the reduction in the levels of OXPHOS complexes containing mtDNA-encoded subunits (Extended Data Fig. 5b). We also observed an increase in the levels of many OXPHOS complex assembly factors (Extended Data Fig. 5c), consistent with a compensatory biogenesis effect typically observed in mice with mitochondrial dysfunction^26, 28, 29^. In addition, we found that levels of most of the mitochondrial ribosomal proteins were drastically decreased (Extended Data Fig. 6a), which is expected as IMT treatment impaired the transcription of the mtDNA-encoded 12S and 16S rRNA (Extended Data Fig. 6b) necessary for the assembly of the small and large subunit of the mitochondrial ribosome^28, 30^.

**Fig. 5:**
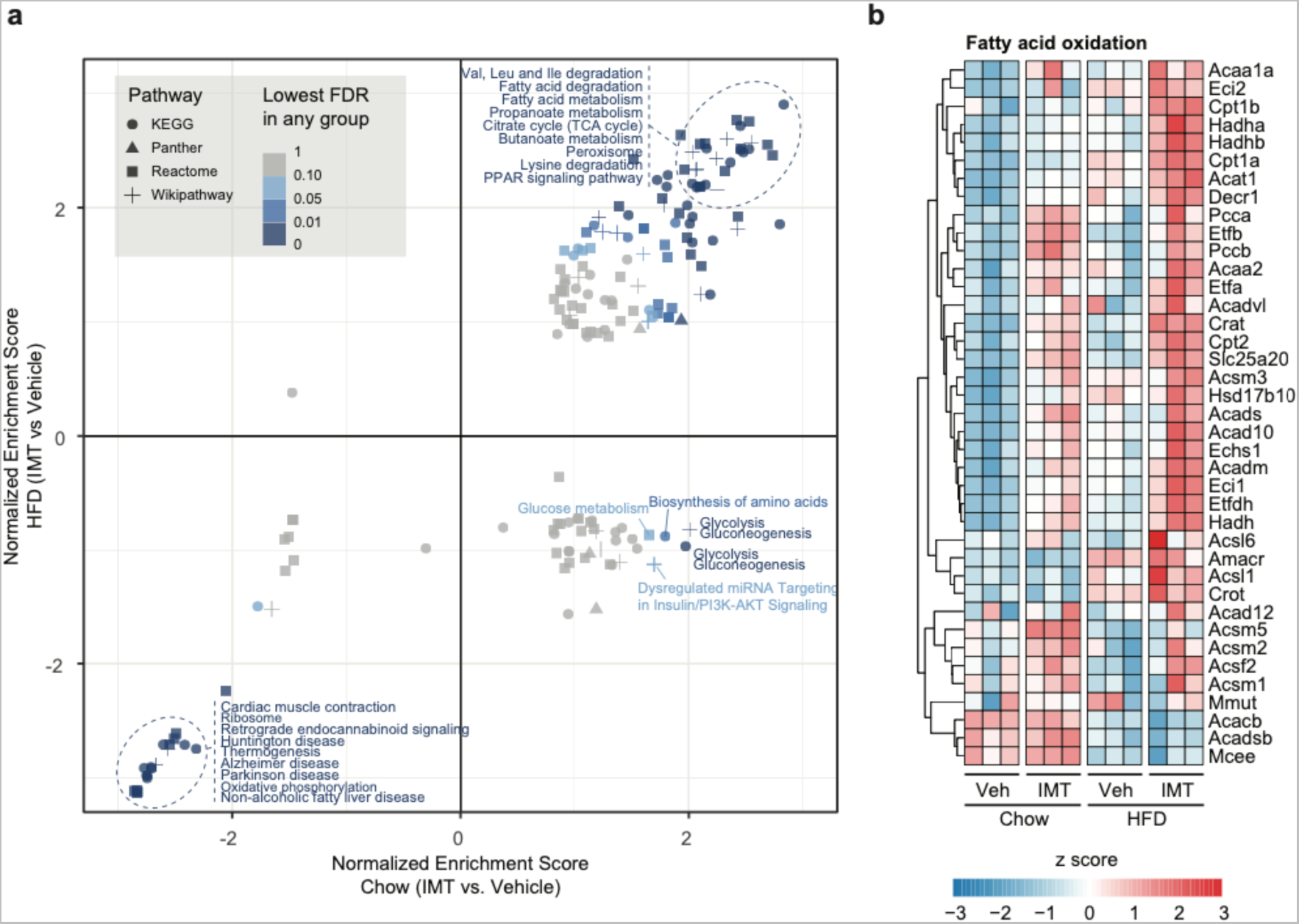
Analysis of the total and mitochondrial proteome. **a**, Gene set enrichment analysis (GSEA) of total tissue and mitochondrial proteomes from liver. **b**, Heatmaps illustrating the protein density of enzymes involved in fatty acid oxidation in mouse livers after four weeks of vehicle or IMT treatment. n = 3 mice per group.

**Fig. 6:**
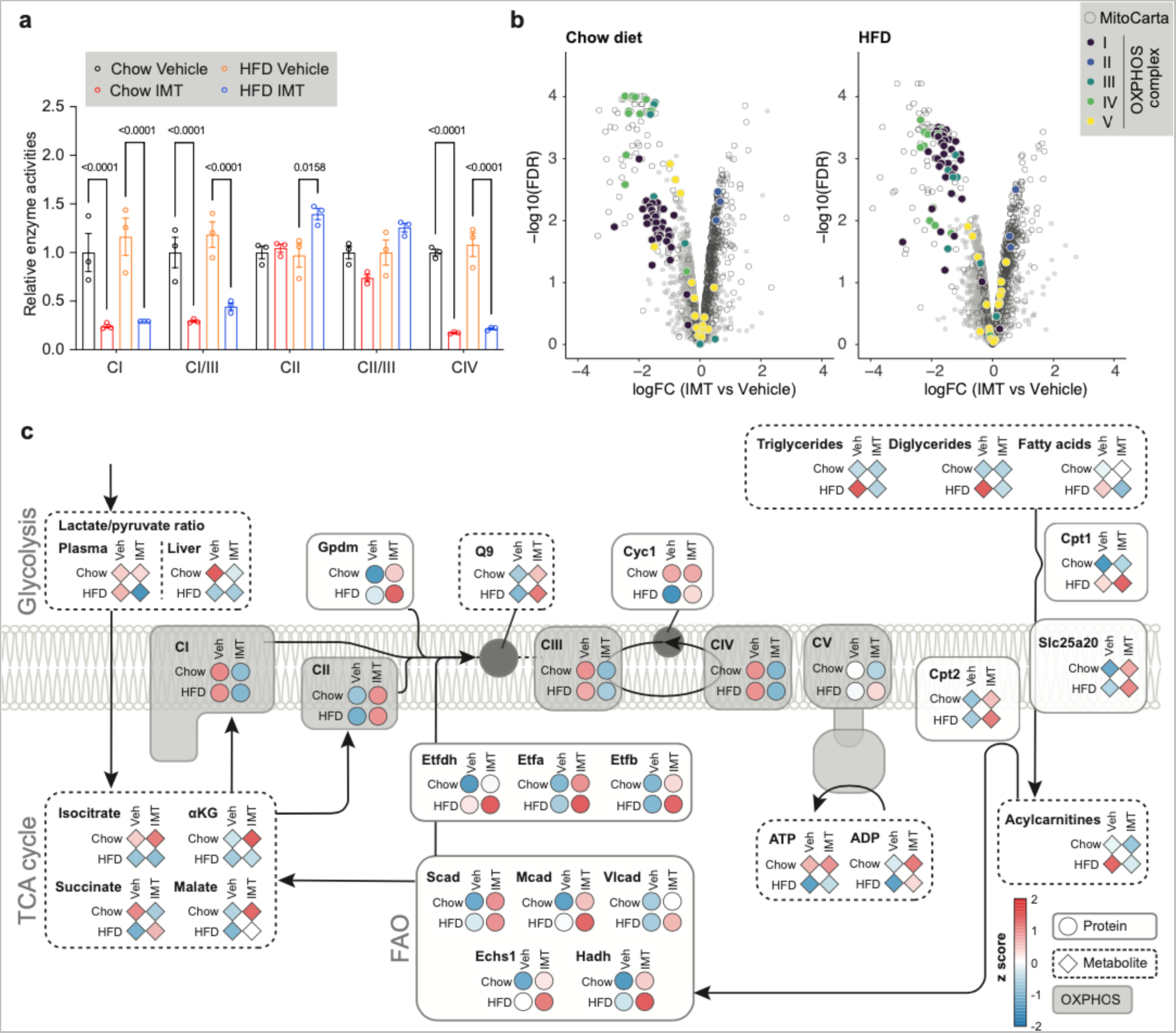
IMT treatment rewires OXPHOS to promote fatty acid oxidation. **a**, Respiratory chain complex activities normalized to citrate synthase activity in liver mitochondria after four weeks of vehicle or IMT treatment. n = 3 mice per group. The analyzed enzyme activities are NADH coenzyme Q reductase (complex I, CI), NADH cytochrome c reductase (complex I/III, CI/III), succinate dehydrogenase (complex II, CII), and cytochrome c oxidase (complex IV, CIV). Data are presented as mean ± SEM. Statistical significance was assessed by a two-way ANOVA with Tukey’s test for multiple comparisons. P-values are shown in the figure. **b**, Changes of all quantified proteins in liver of mice on chow diet or HFD and subjected to vehicle or IMT treatement. Proteins upregulated upon IMT treatment are on the right side of the volcano plot. **c,** An integrated view of the changes of metabolite and protein levels in liver of mice on chow diet (Chow) or HFD and subjected to vehicle (Veh) or IMT treatement.

To further explore the pathways influenced by IMT treatment, we performed gene set enrichment analysis (GSEA) of the total liver tissue proteome and found that enzymes involved in fatty acid metabolism/degradation were markedly enriched in liver of IMT-treated mice, regardless of the diet (Fig. 5a). The GSEA findings indicate that IMT rewires liver metabolism to favor fatty acid degradation, as documented by the increased levels of several key fatty acid oxidation enzymes, e.g., the heterodimeric electron transfer flavoprotein subunits (ETFA and ETFB), electron transfer flavoprotein dehydrogenase (ETF-DH) and the carnitine acyltransferases (CPT1a and CPT2) (Fig. 5b). These findings explain the reduced fat content and increased fat metabolism in whole animals on HFD following IMT treatment (Fig. 1c, e).

To assess OXPHOS function, we measured the activity of respiratory chain enzymes in liver mitochondria and found that IMT treatment caused a marked decrease in the activity of complex I, I/III and IV, whereas the complex II and complex II/III activities were maintained (Fig. 6a). Complex I, III and IV contain critical mtDNA-encoded subunits^31^ and are therefore sensitive to IMT treatment, whereas complex II is exclusively nucleus-encoded and therefore can maintain or even increase its activity (Fig. 6a). The levels of quinones (Q9 and Q10), that receive electrons from different dehydrogenases (e.g., complex I, complex II and ETF-DH) and translocate them to complex III, were normal or increased (Extended Data Fig. 7a). Importantly, the complex III activity that remains after IMT treatment is apparently sufficient to sustain a normal activity of complex II/III (Fig. 6a), despite the marked loss of CIII subunits (Fig. 6b, Extended Data Fig. 5a). The enzyme activity measurements thus show that other dehydrogenases, besides complex I, can sustain normal activity even if the downstream electron translocators, i.e., complex III and IV, have much reduced enzyme levels and activities.

When human cancer cell lines are treated with IMT1B, there is a time-dependent, marked decrease of triphosphate nucleotides (accompanied by increase of mono- and diphosphate nucleotides) and decrease of amino acids, which leads to a cellular energy crisis and cell death^11^. In contrast, metabolite analysis in liver of IMT-treated mice on chow diet or HFD showed normal levels of key mono-, di-, and triphosphate nucleotides (Extended Data Fig. 7b) and amino acids (Extended Data Fig. 7c), consistent with the observation that liver function is not impaired by IMT treatment (Fig. 3e-g).

An integrated view of the changes of metabolite and protein levels in liver shows that IMT treatment causes a massive reprogramming of metabolism to promote fatty acid oxidation regardless of the diet (Fig. 6c).

## Discussion

### A novel principle for drug treatment of obesity

We show here that IMT treatment leads to a dramatic improvement of pathological changes in mice on HFD with normalization of body weight, reversal of hepatosteatosis and reversal of the prediabetic state. This effect is mediated by major reprogramming of metabolism in the liver, caused by activation of fatty acid oxidation pathways (Fig. 6c). Consistent with previous results^11^, IMT treatment markedly impairs the activity of complex I, III and IV, which all contain mtDNA-encoded subunits critical for their function^32^. The OXPHOS system integrates many metabolic pathways through several different dehydrogenases that directly deliver electrons to the Q pool for subsequent electron translocation by complex III and IV and finally reduction of molecular oxygen to water^33^. Although the biogenesis of complex I is critically dependent on mtDNA expression, the other dehydrogenases that directly deliver electrons to the Q pool are exclusively nucleus-encoded and are therefore not directly impacted by IMT treatment (Fig. 6c). The final step of fatty acid oxidation depend on the ETF dehydrogenase that directly delivers electrons to the Q pool for transfer to complex III and IV. Complex I is the main dehydrogenase in the OXPHOS system, and a reduction of its levels and activity may allow better access for ETF dehydrogenase to directly deliver electrons to the Q pool. This type of competition between different dehydrogenases for electron delivery to the Q pool has been documented in budding yeast^34^, and may also be operational in mammalian mitochondria. In summary, we present preclinical evidence for a novel principle for drug treatment of obesity based on inhibition of mtDNA expression in liver to promote fatty acid oxidation. This treatment increases energy expenditure and reduces body weight independent of changes in food intake and physical activity.

## Methods

### IMT substance and vehicle for daily gavage

The IMT substance used in this study (LDC4587) is closely related to the previously published IMT1B^11^. The IMT compound was suspended in vehicle of 0.5% (w/v) hydroxypropyl methylcellulose (Hypromellose, Sigmal Aldrich H3785). Mice were weighed and a volume corresponding to an IMT dose of 30 mg/kg body weight was given by gavage once per day.

### Pharmacokinetics analyses

LDC4857 was extracted from plasma and tissues by protein precipitation using acetonitrile. Tissue samples were homogenized using two parts (w/v) of PBS prior to extraction with acetonitrile. Following filtration, samples were analyzed by liquid chromatography tandem-mass spectrometry using a Prominence UFLC system (Shimadzu) coupled to a Qtrap 5500 instrument (ABSciex). Test articles were separated on a C18 column using a gradient elution with an acetonitrile/water mixture containing 0.1% formic acid as the mobile phase. Chromatographic conditions and mass spectrometry parameters were optimized for the tested compound prior to sample analysis. Concentrations of LDC4857 were calculated by means of a standard curve.

### Mouse models

Male mice (C57BL/6N) were purchased from Charles River Laboratories (Germany) at four weeks of age. Mice were maintained under a 12 hours light/dark cycle and had free access to water and standard chow diet (4% kcal from fat; Special Diet Service, UK) or a high-fat diet (60% kcal from fat, TD.06414; Envigo) ad libitum for 12 weeks. After eight weeks on chow diet or HFD, mice received daily oral gavage of IMT compound or vehicle for a continuous 28 days. Animal studies were approved by the animal welfare ethics committee and performed in compliance with National and European law.

### Indirect calorimetry

The Oxymax/Comprehensive Lab Animal Monitoring System (CLAMS, Columbus, U.S.A) cage system was used to measure food intake, oxygen consumption, carbon dioxide production, energy expenditure, respiratory exchange ratio (RER) and physical activity. On the first day, mice were weighed and their overall condition were checked before placing them individually into the metabolic cage. After 24-48 hours acclimation (day three), mice were visually checked for signs of stress and food/drink consumption was recorded from 09:00. If mice were not eating and/or drinking (determined as less than 1 g of food or 1 mL of water), they were removed from the experiment (n = 1 mouse was excluded). From 18:00 of day three to the 12:00 of day five the data collection phase was performed. From 18:00 of day four to the 6:00 of day five, food was removed from the cage and mice were underwent a 12 hour fasting period. At 6:00 of day five, food was given to the mice and data were collected for the refeeding phase. Caloric consumption was calculated using the following values: HFD 5.24 kcal/g, chow diet 3.18 kcal/g.

### Intraperitoneal glucose tolerance test

Experiments were performed following 4 hours of fasting starting at ∼06:00. Blood glucose was monitored using the Contour XT glucometer (Bayer) from samples collected at the distal tail vein. Following an initial blood glucose measurement, glucose (2 g/kg body weight) was injected intraperitoneally. Blood glucose was measured 15, 30, 60, 90, and 120 min after the injection.

### *Ex vivo* glucose stimulated insulin secretion and insulin measurement

Mouse islets were isolated as previously described^35^. Handpicked islets were incubated in RPMI medium (11875093, Thermo Fisher Scientific) with 11 mM glucose, 10% FBS and 1% Penicillin-Streptomycin overnight to recover from the isolation. Glucose-stimulated insulin secretion was performed as previously described^36^. Briefly, 15 islets from each group were equilibrated in KRBH solution containing 2.8 mM glucose for 2 hours, then transferred to be incubated in KRBH containing 2.8 mM glucose for 1 hour, 16.7 mM glucose for another hour, and the supernatant from each incubation was collected. Islets were lysed with RIPA (R0278, Sigma) containing protease inhibitor cocktail (11697498001, Roche) and phosphatase inhibitor cocktail (4906845001, Roche) to determine protein concentration. The measurement of insulin in the islet lysate, the supernatant and the fasting and 15 min GTT serum was performed using Rat/Mouse Insulin ELISA kit (EZRMI-13K, Sigma-Aldrich).

### Histology

Epididymis white adipose tissue (eWAT) and one liver lobe were collected and fixed in 4% paraformaldehyde at 4°C for 24 hours. The tissues were processed routinely and embedded in paraffin and sectioned to 5 μm thickness. Hematoxylin & eosin (H&E) staining was performed to analyze morphology of the tissues.

### Hepatic lipid quantification

Quantification of liver triglycerides was performed using a triglyceride quantification kit (ab65336, Abcam) according to the manufacturer’s instructions.

### Crude mitochondrial isolation

Mitochondria from liver were isolated by differential centrifugation in mitochondrial isolation buffer (320 mM Sucrose, 10 mM Tris-HCl, pH 7.4, 1 mM EDTA and 0.2% BSA), supplemented with EDTA-free complete protease inhibitor cocktail and PhosSTOP Tablets (Roche). Liver tissue was homogenized using a Potter homogenizer on ice (13 strokes, 500 rpm). Nuclei and cell debris were pelleted at 1,000 g/ 10 min/ 4°C. Mitochondria were pelleted from the supernatant by centrifugation at 10,000 g/ 10 min/ 4°C. The mitochondrial pellet was carefully resuspended in mitochondrial isolation buffer without BSA and the differential centrifugations were repeated to obtain crude mitochondria.

### Biochemical evaluation of respiratory chain enzyme activity

The measurement of respiratory chain enzyme activities and citrate synthase activity was performed as previously described^37^.

### Western blot analysis

Mitochondrial proteins (10 μg) were resuspended in 1× NuPAGE™ LDS Sample Buffer. Mitochondrial proteins were thereafter separated by SDS–PAGE (4–12% Bis-Tris Protein Gels; Invitrogen) and transferred onto polyvinylidene difluoride (PVDF) membranes (Merck Millipore). Immunoblotting was performed using standard procedures with ECL reagent detection.

### RNA extraction and RT-qPCR

RNA was extracted from mouse liver, skeletal muscle, eWAT, heart and BAT using Trizol reagent (Invitrogen) according to the manufacturer’s instructions and then treated with TURBO DNA-free™ DNase (Invitrogen). For RT-qPCR expression analysis, cDNA was reversed transcribed from 1 μg total RNA using the High-Capacity cDNA Reverse

Transcription Kit (Invitrogen). The qPCR was performed in a QuantStudio 6 Flex Real-Time PCR System (Life Technologies), using TaqMan™ Universal Master Mix II, with UNG (Applied Biosystems) to quantify mitochondrial transcripts (mt-rRNAs and mt-mRNAs), actin, and 18S rRNA.

### DNA isolation and mtDNA quantification

Genomic DNA was isolated from mouse liver, eWAT and skeletal muscle using the DNeasy Blood and Tissue Kit (Qiagen) following manufacturer’s instructions and treated with RNase A. Levels of mtDNA were measured by quantitative PCR using 5 ng of DNA in a QuantStudio 6 Flex Real-Time PCR System using TaqMan™ Universal Master Mix II, with UNG. Nd1, Cox1, and Cyb probes were used for TaqMan assays to measure mtDNA levels. 18S was used for normalization.

### Isolation of ultrapure mitochondria

Ultrapure mitochondria were prepared as previously described^30^. In brief, crude mitochondrial pellets from mouse liver were washed in 1xM buffer (220 mM mannitol, 70 mM sucrose, 5 mM HEPES pH 7.4, 1 mM EGTA pH 7.4); pH was adjusted with potassium hydroxide, supplemented with EDTA-free complete protease inhibitor cocktail and PhosSTOP Tablets (Roche), and purified on a Percoll density gradient (12%:19%:40%) via centrifugation in a SW41 rotor at 42,000 g at 4 °C for 1 hour in a Beckman Coulter Optima L-100 XP ultracentrifuge using 14 × 89 mm Ultra-Clear Centrifuge Tubes (Beckman Instruments Inc.). Mitochondria were harvested at the interphase between 19 and 40% Percoll, and washed three times with buffer 1xM, and mitochondrial pellets were frozen at −80 °C.

### Label-free quantitative proteomics

#### Proteomics sample preparation

Frozen tissue pieces were placed in precooled “Lysing Matrix D” tubes, followed by addition of 400 µL lysis buffer (1 % SDC in 100 mM Tris-HCl, pH 8.5). Tissue pieces were lysed at 4

°C by 3 cycles of 40 s bead beating (6.0 setting) and 20 s pause in the FastPrep-24 (MP Biomedicals). Thereafter, lysates were transferred into reaction tubes and boiled for 10 min at 95 °C. Similarly, ultrapure mitochondria pellets were resuspended in 150 µL lysis buffer and boiled for 10 min at 95 °C. After lysate boiling, the protein concentration was estimated by tryptophan assay and 30 µg of each sample were diluted with lysis buffer to a protein concentration of 0.75 µg/µL. Proteins were reduced and alkylated by adding chloroacetamide (CAA) and Tris(2-carboxyethyl)phosphine (TCEP) to a final concentration of 40 mM and 10 mM, respectively, and 5 min incubation at 45 °C proteins. After adding Trypsin (1:100 w/w, Sigma-Aldrich) and LysC (1/100 w/w, Wako), proteins were digested overnight at 37 °C. Protein digestion was quenched by adding 200 µL 1% TFA in isopropanol to the samples. Subsequently, peptides were loaded onto SDB-RPS StageTips (Empore) followed by washes with 200 µL 1% TFA in isopropanol and 200 µL 0.2% TFA in 2% ACN. Peptides were eluted with 60 µL 1.25% NH_4_OH in 80% ACN and dried in a SpeedVac centrifuge (Eppendorf, Concentrator plus). Dried peptides were resuspended in A* (0.2% TFA in 2% ACN) and subjected to measurement by LC-MS/MS.

#### LC-MS/MS and proteomics data analysis

Peptide concentration was estimated by NanoDrop and 250 ng peptide material were used for individual measurements. Peptides were loaded onto a 50 cm, in-house packed, reversed-phase columns (75 μm inner diameter, ReproSil-Pur C18-AQ 1.9 μm resin [Dr. Maisch GmbH]) and separated with a binary buffer system consisting of buffer A (0.1% FA) and buffer B (0.1% FA in 80% ACN) with an EASY-NLC 1,200 (Thermo Fisher Scientific). The LC system was directly coupled online with the mass spectrometer (Exploris 480, Thermo Fisher Scientific) via a nano-electrospray source. Peptide separation was performed at a flowrate of 300 µL/min and an elution gradient starting at 5% B increasing to 30% B in 80 min, 60% in 4 min and 95% in 4 min.

Data were acquired in DIA mode with a scan range of 300-1650 m/z at a resolution of 120,000. The AGC was set to 3e6 at a maximum injection time of 60 ms. Precursor fragmentation was achieved via HCD (NCD 25.5%, 27.5%, 30%) and fragment ions were analyzed in 33 DIA windows at a resolution of 30,000, while the AGC was kept at 1e6.

DIA raw files were processed using Spectronaut (v14) with default settings. Perseus (v1.6.7.0)^38^ was used on data with three valid values in at least one treatment group. Principal component analyses were performed on missing-values imputed matrices, and ANOVA testing with permutation-based FDR correction. Gene Set Enrichment Analyses were computed with WebGestalt 2019^39^ in an R environment (v4.1.2) correcting for multiple library testing, and normalized enrichment scores were reported. Statistical analyses and FDR calculations were performed with limma, and an FDR cutoff of 0.05 was defined as significant. Heatmaps were generated on filtered, imputed and z-transformed data matrices in R with the pheatmap package.

### Metabolomics and lipidomics

#### Samples extraction of polar and lipid metabolites

Metabolites were extracted from 10-15 mg of frozen tissue. The frozen tissue samples were homogenized to a fine tissue powder, using a ball mill (MM400, Retsch, Haan, Germany).

After the tissue was pulverized in 2 mL round-bottom microcentrifuge tubes, 1 mL of a -20 °C methyl-tert butyl-ether:methanol:water (5:3:2 (v:v:v)) mixture, containing 0.2 µL/mL of the deuterated EquiSplash lipidomix (Avanti, Birmingham, AL, USA), 0.2 µL/mL of the U-13C15N amino acid mix (Cambridge isotopes MSK_A2-1.2), 0.1 uL/mL of 1 mg/mL 13C10 ATP, 15N5 ADP and 13C1015N5 AMP (Sigma) and 0.2 uL/mL 100 µg/m of deuterated citric acid, as Internal standards, was added to each sample. After addition of the extraction buffer, the samples were immediately vortexed before they were incubated for additional 30 min at 4 °C on an orbital shaker. Proteins were removed by a 10 min 21,000 g centrifugation at 4 °C and the supernatant was transferred to a fresh 2 mL Eppendorf tube. To separate the organic from the polar phase, 150 μL of MTBE and 100 µL of UPC/MS-grade water was added to the cleared supernatant, which was shortly vortexed before mixing it for 15 min at 15 °C on an orbital shaker. Phase separation was obtained after a 5 min centrifugation at 16,000 g at 15 °C. The upper MTBE phase, which contains the hydrophobic compounds (lipids), was sampled to a fresh 1.5 mL microcentrifuge tube (∼600 µL), while the remaining polar phase (∼600 µL) was kept in the initial 2 mL tube. These two fractions were then immediately concentrated to dryness in a speed vacuum concentrator (LaboGene, MaxiVac, Allerod, Denmark) at room temperature. Thereafter, samples were then either stored at -80 °C or processed immediately for LC-MS analysis.

#### Anion-Exchange Chromatography Mass Spectrometry (AEX-MS) for the analysis of anionic metabolites

The polar phase of the extracted metabolites was re-suspended in 400 µL of ULC/MS-grade water (Biosove, Valkenswaard, Netherlands). After 15 min of incubation on a thermomixer at 4 °C and a 5 min centrifugation at 16,000 g at 4 °C, 100 µL of the cleared supernatant were transferred to polypropylene autosampler vials (Chromatography Accessories Trott, Germany).

The samples were analyzed using a Dionex ionchromatography system (Integrion, Thermo Fisher Scientific) as described previously. In brief, 5 µL of polar metabolite extracts were injected in full loop mode using an overfill factor of 1, onto a Dionex IonPac AS11-HC column (2 mm × 250 mm, 4 μm particle size, Thermo Fisher Scientific) equipped with a Dionex IonPac AG11-HC guard column (2 mm × 50 mm, 4 μm, Thermo Fisher Scientific). The column temperature was held at 30 °C, while the auto sampler was set to 6 °C. A potassium hydroxide gradient was generated using a potassium hydroxide cartridge (Eluent Generator, Thermo Scientific), which was supplied with deionized water. The metabolite separation was carried at a flow rate of 380 µL/min, applying the following gradient conditions: 0-3 min, 10 mM KOH; 3-12 min, 10-50 mM KOH; 12-19 min, 50-100 mM KOH, 19-21 min, 100 mM KOH, 21-22 min, 100-10 mM KOH. The column was re-equilibrated at 10 mM for 8 min.

For the analysis of metabolic pool sizes the eluting compounds were detected in negative ion mode using full scan measurements in the mass range m/z 50–750 on a Q-Exactive HF high resolution MS (Thermo Fisher Scientific). The heated electrospray ionization (ESI) source settings of the mass spectrometer were: Spray voltage 3.2 kV, capillary temperature was set to 300 °C, sheath gas flow 50 AU, aux gas flow 20 AU at a temperature of 330 °C and a sweep gas flow of 2 AU, while the S-lens was set to a value of 60 AU. The semi-targeted LC-MS data analysis was performed using the TraceFinder software (Version 4.1, Thermo Fisher Scientific). The identity of each compound was validated by authentic reference compounds, which were measured at the beginning and the end of the sequence. For data analysis, the area of the deprotonated [M-H+]-monoisotopic mass peak of each compound was extracted and integrated using a mass accuracy <5 ppm and a retention time (RT) tolerance of <0.05 min as compared to the independently measured reference compounds. Areas of the cellular pool sizes were normalized to the internal standards, which were added to the extraction buffer, followed by a normalization to the fresh weight of the analyzed sample.

#### Semi-targeted liquid chromatography-high-resolution mass spectrometry-based (LC-HRS-MS) analysis of amine-containing metabolites

The LC-HRMS analysis of amine-containing compounds was performed as described previously^11^. In brief, 50 µL of the resuspended 400 µL of the above mentioned (AEX-MS section) polar phase, were mixed with 25 µL of 100 mM sodium carbonate (Sigma), followed by the addition of 25 µL 2% [v/v] benzoylchloride (Sigma) in acetonitrile (UPC/MS-grade, Biosove, Valkenswaard, Netherlands). Derivatized samples were thoroughly mixed and transferred to autosampler vials, where they were kept at 20 °C until analysis.

For the LC-MS analysis, 1 µL of the derivatized sample was injected onto a 100 x 2.1 mm HSS T3 UPLC column (Waters). The flow rate was set to 400 µL/min using a binary buffer system consisting of buffer A (10 mM ammonium formate (Sigma), 0.15% [v/v] formic acid (Sigma) in UPC-MS-grade water (Biosove, Valkenswaard, Netherlands). Buffer B consisted of acetonitrile (IPC-MS grade, Biosove, Valkenswaard, Netherlands). The column temperature was set to 40 °C, while the LC gradient was: 0% B at 0 min, 0-15% B 0-4.1 min; 15-17% B 4.1– 4.5 min; 17-55% B 4.5-11 min; 55-70% B 11 – 11.5 min, 70-100% B 11.5 - 13 min; B 100% 13 - 14 min; 100-0% B 14 -14.1 min; 0% B 14.1-19 min; 0% B. The mass spectrometer (Q-Exactive Plus, Thermo Fisher Scientific) was operating in positive ionization mode recording the mass range m/z 100-1000. The heated ESI source settings of the mass spectrometer were: Spray voltage 3.5 kV, capillary temperature 300 °C, sheath gas flow 60 AU, aux gas flow 20 AU at 330 °C and the sweep gas was set to 2 AU. The RF-lens was set to a value of 60.

Semi-targeted data analysis for the samples was performed using the TraceFinder software (Version 4.1, Thermo Fisher Scientific). The identity of each compound was validated by authentic reference compounds, which were run before and after every sequence. Peak areas of [M + nBz + H]+ ions were extracted using a mass accuracy (<5 ppm) and a retention time tolerance of <0.05 min. Areas of the cellular pool sizes were normalized to the internal standards (U-15N;U-13C amino acid mix (MSK-A2-1.2), Cambridge Isotope Laboratories), which were added to the extraction buffer, followed by a normalization to the fresh weight of the analyzed sample.

#### Liquid Chromatography-High Resolution Mass Spectrometry-based (LC-HRMS) analysis of lipids

The dried lipid fractions were resuspended in 400 µL of UPLC-grade acetonitrile: isopropanol (70:30 [v:v], Biosove, Valkenswaard, Netherlands). Samples were vortexed for 10 s and incubated for 10 min on a thermomixer at 4 °C. Resuspended samples were centrifuged for 5 min at 10,000 g and 4 °C, before transferring the cleared supernatant to 2 mL glass vials with 200 µL glass inserts (Chromatography Zubehör Trott, Germany). All samples were placed in a UHPLC sample manager (Vanquish, Thermo Fisher Scientific, Bremen, Germany), which was set to 6 °C. The UHPLC was connected to a Tribrid Orbitrap HRMS, equipped with a heated ESI (HESI) source (ID-X, Thermo Fischer Scientific).

A volume of 1 µL of each lipid sample was injected into a 100 x 2.1 mm BEH C8 UPLC column, packed with 1.7 µm particles (Waters). The flow rate of the UPLC was set to 400 µL/min and the buffer system consisted of buffer A (10 mM ammonium acetate, 0.1% acetic acid in UPLC-grade water) and buffer B (10 mM ammonium acetate, 0.1% acetic acid in UPLC-grade acetonitrile/isopropanol 7:3 [v/v]). The UPLC gradient was as follows: 0-1 min 45% A, 1-4 min 45-25% A, 4-12 min 25-11% A, 12-15 min 11-1% A, 15-20 min 1% A, 20-20.1 min 1-45% A and 20.1-24 min re-equilibrating at 45% A, which leads to a total runtime of 24 min per sample and polarity.

The ID-X mass spectrometer was operating for the first injection in positive ionization mode and for the second injection in negative ionization mode. In both cases, the analyzed mass range was between m/z 150-1500. The resolution was set to 120,000, leading to approximately 4 scans per second. The RF lens was set to 50%, while the AGC target was set to 100%. The maximal ion time was set to 100 ms and the HESI source was operating with a spray voltage of 3.6 kV in positive ionization mode, while 3.2 kV were applied in negative ionization mode. The ion tube transfer capillary temperature was 300 °C, the sheath gas flow 60 arbitrary units (AU), the auxiliary gas flow 20 AU and the sweep gas flow was set to 1 AU at 330 °C.

To obtain positive ionization-mode and negative ionization-mode MS/MS-based lipid annotations, we performed five iterative MS/MS deep sequencing runs on a pooled sample of each tissue type using the AcquireX algorithm (Xcalibur Version 4.3, Thermo Fisher Scientific). Lipids from these MS/MS spectra were then automatically annotated using LipidSearch (Version 4.2, Thermo Fisher Scientific). The annotated lipids were filtered for quality grades (A,B and C were accepted) and the resulting lipid IDs, m/z and retention time values were exported into a TraceFinder compound data base (Version 4.1, Thermo Fisher Scientific). From this compound database we generated a TraceFinder method for each sample set and extracted the corresponding peaks from each full scan MS spectrum.

For data analysis the area of each monoisotopic mass peak was extracted and integrated using a mass accuracy <5 ppm and a retention time (RT) tolerance of <0.05 min as compared to the independently measured reference compounds. Areas of the cellular pool sizes were normalized to the internal standards, followed by a normalization to the fresh weight/volume of the analyzed sample.

### Statistical analysis

Experiments were performed at least three times, and results are representative of n > 5 independent biological replicates, unless indicated otherwise. All values are presented as mean

± SEM. Statistical analyses were conducted using the GraphPad Prism software (v.9.4.0). Statistical significance was assessed by a two-way ANOVA with Tukey’s test for multiple comparisons. P-values are shown in the figure.

## Acknowledgements

We thank the Morphological Phenotype Analysis Core Facility (FENO) at Karolinska Institutet for assistance with histology and imaging. We thank M. Moedas and G. Gao for technical assistance. Figure 1a was created using BioRender.com. **Funding:** NGL was supported by the Swedish Research Council (2015-00418), Swedish Cancer Foundation (CAN2018/602), the Knut and Alice Wallenberg foundation, European Research Council (Advanced Grant 2016-741366), The Swedish Diabetes Foundation (DIA2020-516, DIA2021-620), Novo Nordisk Foundation (NNF20OC006316) and grants from the Swedish state under the agreement between the Swedish government and the county councils (RS2020-0731). LSK and FAS were supported by EMBO long-term fellowships (ALTF 570-2019 and ALTF 399-2021). JRZ was supported by Knut and Alice Wallenberg Foundation (2021.0249), Swedish Research Council (2015-00165), and the Strategic Research Programme in Diabetes at Karolinska Institutet (Swedish Research Council 2009-1068).

## Author contributions

NGL and SJ conceived the project, designed the experiments, and wrote the manuscript. SJ and TY performed and interpreted the majority of the experiments. NGL, SJ, LSK, CK, OA, PG, AK, JRZ and MMann advised on methodology. LSK, DRG, MMennuni, RF, DA, JM, MB, LR, and RW performed experiments and analyzed the data. FAS, FMH, LSK and PG performed and interpreted the proteomics and metabolomics experiments. AU, TB, and RDL supervised the inhibitor generation and profiling, established the structure-activity- and property-relationships and developed the proper formulation and dosing regimen for the *in vivo* study. NGL supervised the project. All the authors commented on the manuscript.

## Additional information

None

## Data Availability

The mass spectrometry proteomics data have been deposited to the ProteomeXchange Consortium via the PRIDE partner repository^40^ with the dataset identifier PXD034771.

---

**Extended Data Fig. 1:**
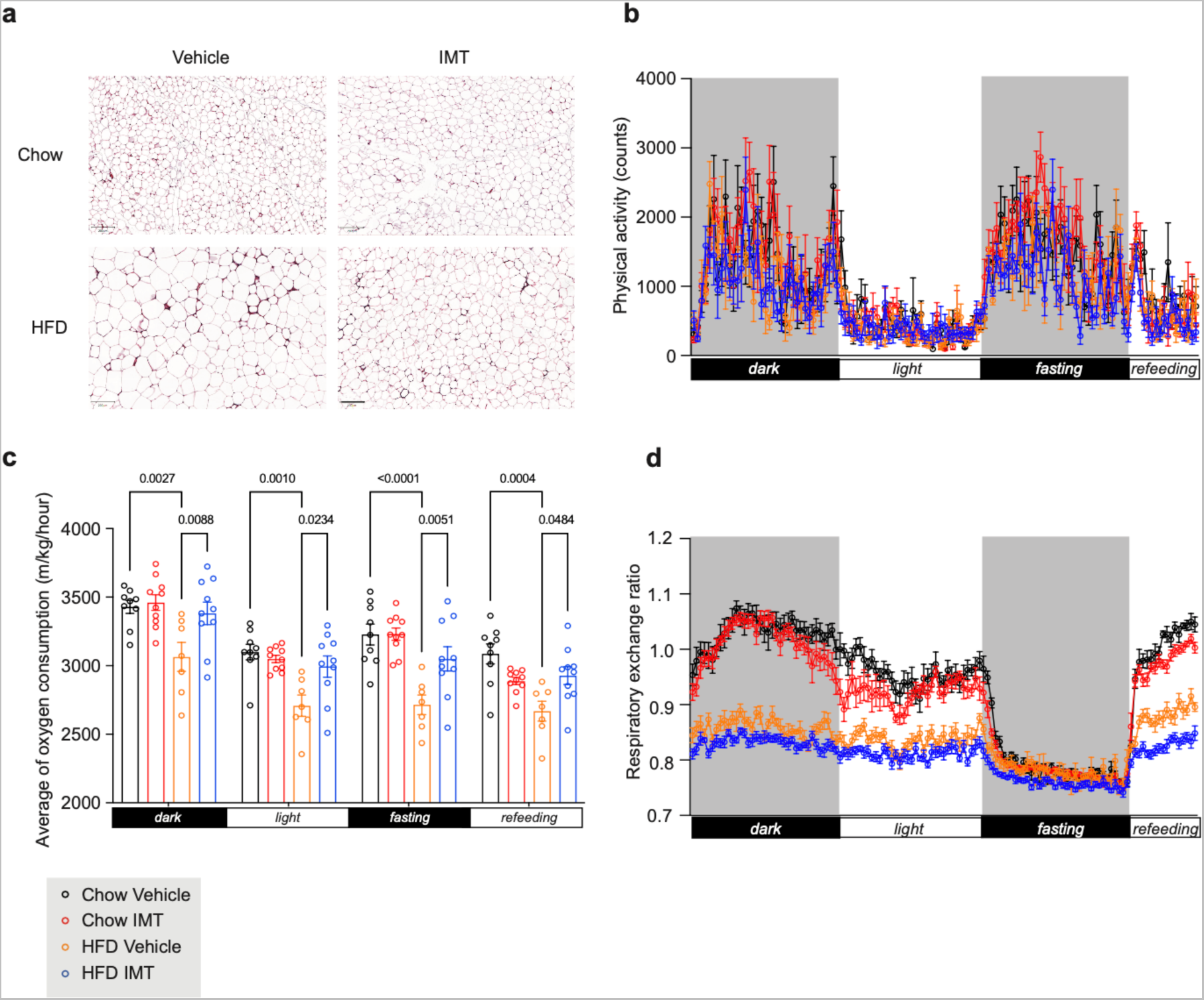
Fat metabolism in mice treated with IMT. **a**, Representative images of H&E staining of eWAT. Scale bar, 200 μm. n = 5 mice per group. **b-d**, Measurement of whole-body metabolism during the fourth week of gavage treatment with vehicle or IMT compound by using the Oxymax/Comprehensive Lab Animal Monitoring System (CLAMS). The first three days were used to acclimate the animals to the CLAMS system, followed by measurements during the fourth day. Day five included a 12 hours period of fasting followed by six hours of refeeding. **b,** Physical activity during day four and day five. **c**, The average oxygen consumption rate during day four and day five. **d**, Respiratory exchange ratio during day four and day five. Chow vehicle n = 10, Chow IMT n = 10, HFD vehicle n = 8, and HFD IMT n = 11 mice. All data are presented as mean ± SEM. Statistical significance was assessed by a two-way ANOVA with Tukey’s test for multiple comparisons. P-values are shown in the figure.

**Extended Data Fig. 2:**
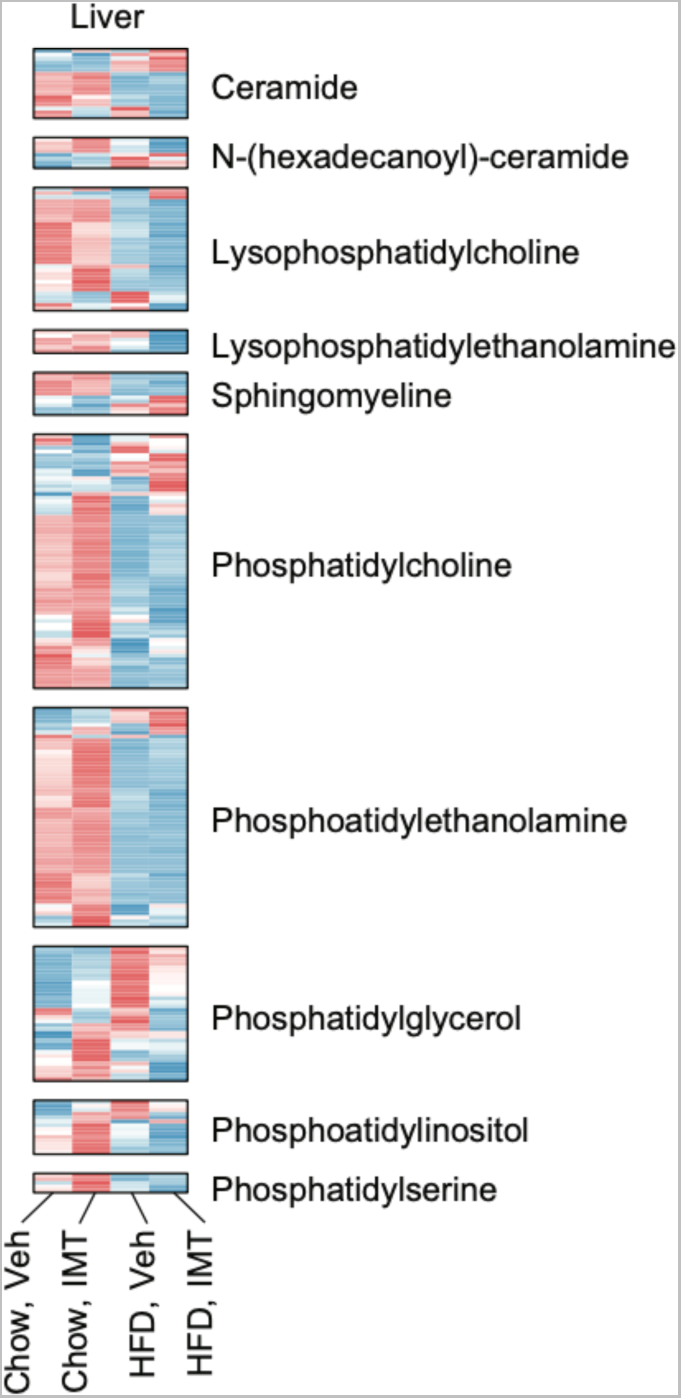
IMT treatment does not change levels of phospholipids and sphingolipids. Levels of phospholipids and sphingolipids in mouse liver after four weeks of vehicle or IMT treatment. n = 8 mice per group.

**Extended Data Fig. 3:**
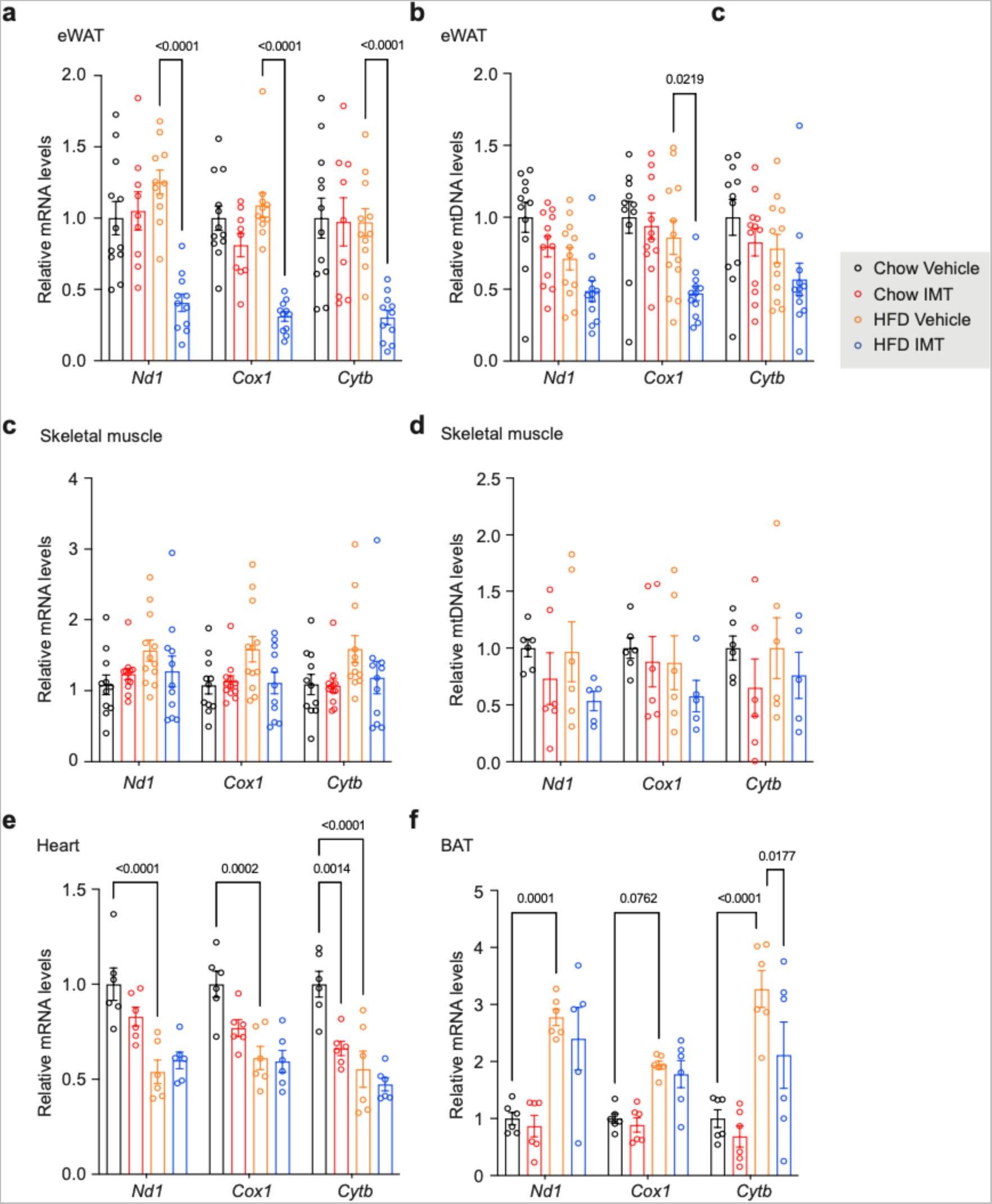
Levels of mitochondrial transcripts and mtDNA in different tissues of IMT-treated mice. **a-f**, Levels of representative mitochondrial transcripts and mtDNA were measured in tissues of mice on chow diet or HFD treated with vehicle or IMT compound for four weeks. The mtDNA transcript (**a**) and mtDNA (**b**) levels in eWAT. n = 6 mice per group. The mtDNA transcript (**c**) and mtDNA (**d**) levels in skeletal muscle. n = 6 mice per group. The mtDNA transcript levels in heart (**e**) and brown adipose tissue (BAT, **f**). n = 6 mice per group. All data are presented as mean ± SEM. Statistical significance was assessed by a two-way ANOVA with Tukey’s test for multiple comparisons. P-values are shown in the figure.

**Extended Data Fig. 4:**
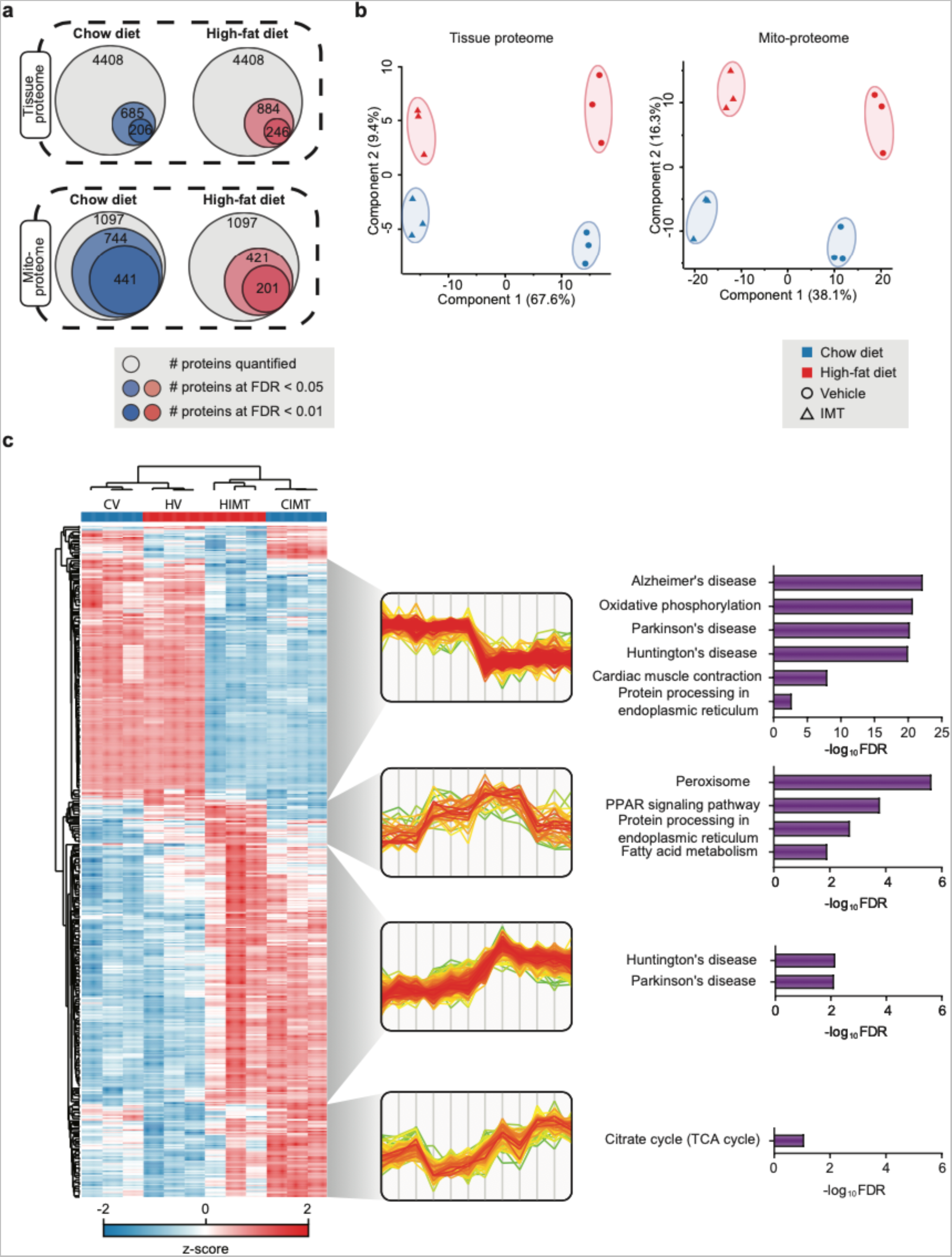
Analysis of the total and mitochondrial proteome. **a**, Venn diagram showing number of quantified and significantly changed proteins at a given FDR cutoff. The liver tissue proteome and the proteome of isolated liver mitochondria (Mitoproteome) are shown. **b,** Principal component analyses (PCA) of the liver tissue and liver mitochondrial proteomes. **c**, Hierarchical clustering analysis of the total proteome (ANOVA-significant proteins at FDR < 5%; left) with biological replicates as individual lanes (CV: control vehicle; HV: HFD vehicle; HIMT: HFD IMT-treated, CIMT: control IMT-treated), z-score normalised fold changes in indicated clusters (middle) with the same sample order as in the heatmap, and significant KEGG terms in each cluster (Fisher exact test at FDR < 5%; right).

**Extended Data Fig. 5:**
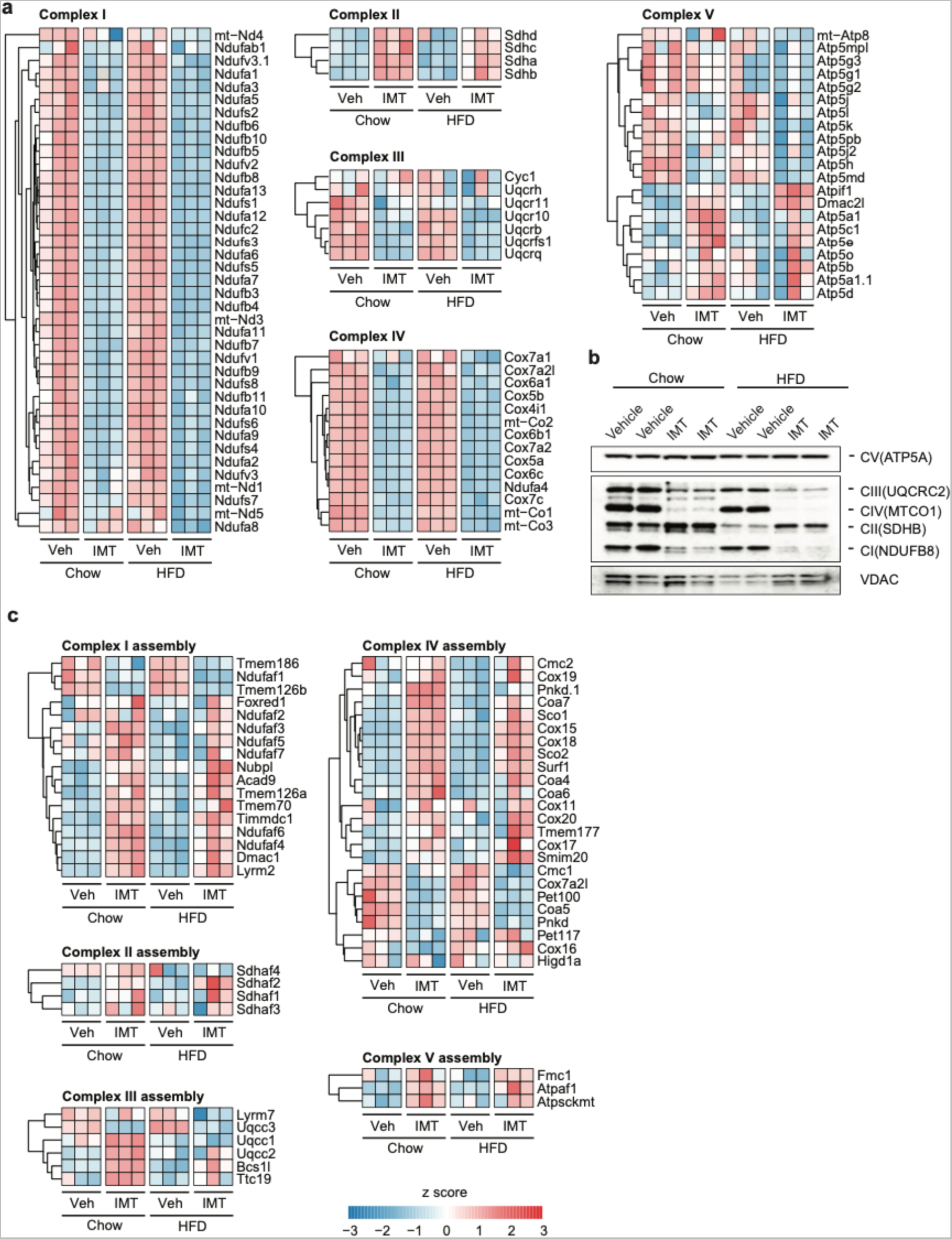
IMT treatment rewires OXPHOS. **a**, Heatmaps illustrating the protein density of subunits of OXPHOS complexes in mouse liver after four weeks of vehicle or IMT treatment. n = 3 mice per group. **b,** Representative western blot analyses of OXPHOS protein levels in liver mitochondria after four weeks of vehicle or IMT treatment. Subunits of complex I (NDUFB8), complex II (SDHB), complex III (UQCRC2), complex IV (MTCOX1), and complex V (ATP5A) were analyzed. VDAC was used as loading control. A representative image of n = 3 independent experiments is shown. **c,** Heatmaps depicting the protein density of different mitochondrial OXPHOS assembly factors in mouse liver after four weeks of vehicle or IMT treatment. n = 3 mice per group.

**Extended Data Fig. 6:**
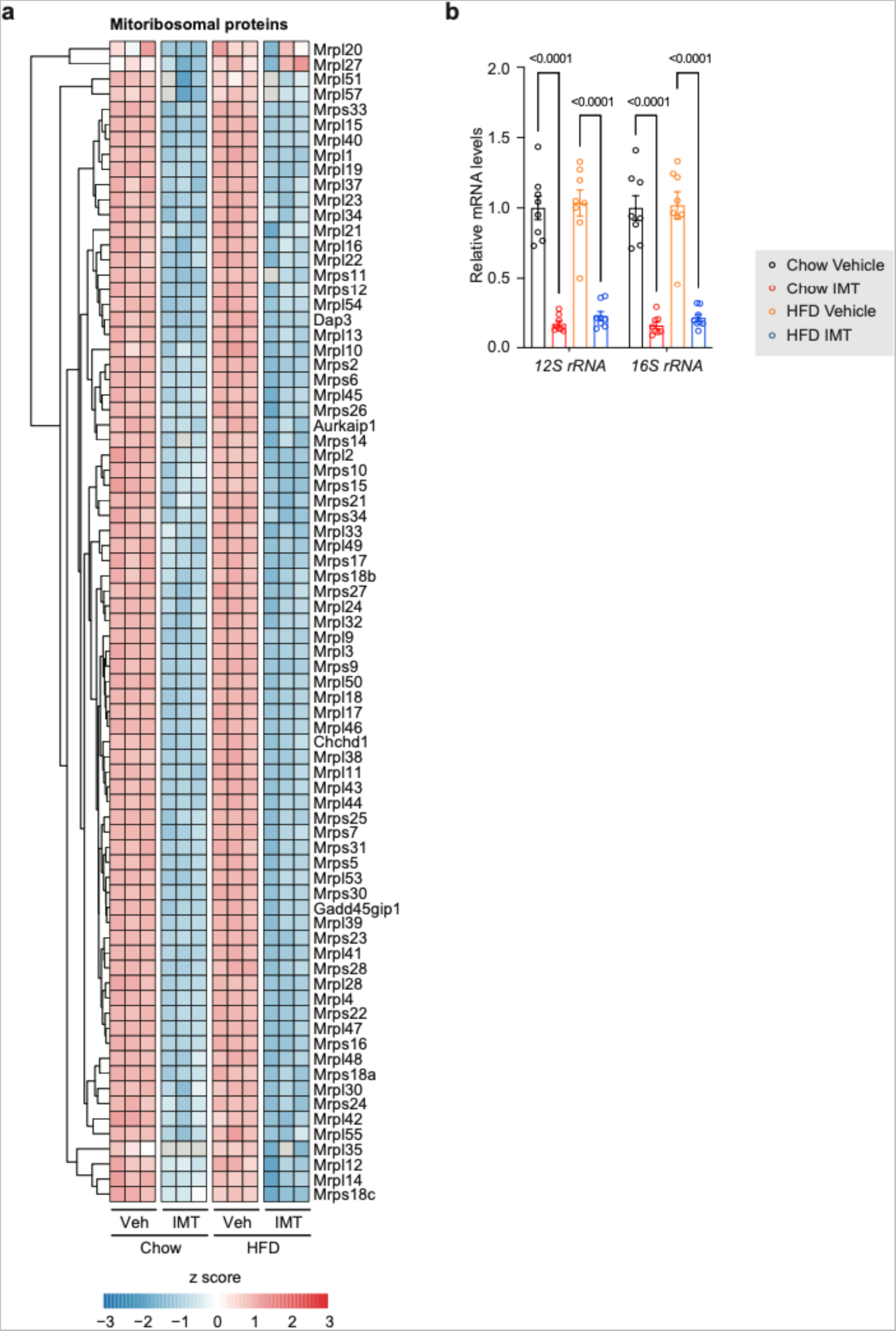
IMT decreases mitoribosomal proteins and rRNAs. **a**, Heatmaps depicting the protein density of mitoribosomal proteins in mouse liver after four weeks of vehicle or IMT treatment. n = 3 mice per group. **b,** The mtDNA-encoded 12S and 16S rRNA transcripts in mouse liver after four weeks of vehicle or IMT treatment. n = 8 mice per group. Data are presented as mean ± SEM. Statistical significance was assessed by a two-way ANOVA with Tukey’s test for multiple comparisons. P-values are shown in the figure.

**Extended Data Fig. 7:**
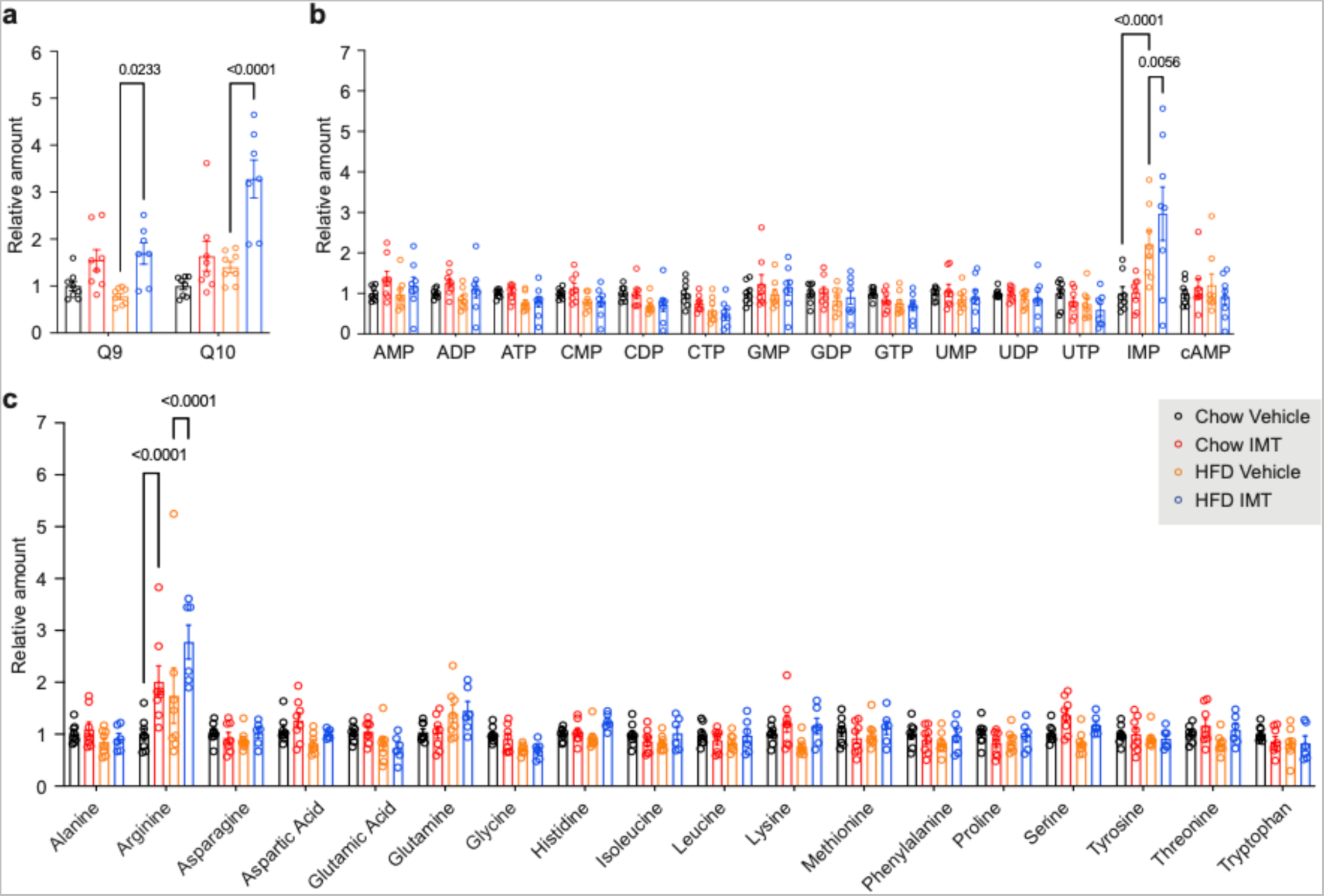
Analysis of the liver metabolites. **a**, Fold change in the levels of quinones (Q9 and Q10) in mouse liver after four weeks of vehicle or IMT treatment. Chow Vehicle, Chow IMT and HFD Vehicle n = 8 mice per group, HFD IMT n = 7 mice per group **b,** Fold change in nucleotide levels in mouse liver after four weeks of vehicle or IMT treatment. n = 8 mice per group. **c**, Fold change in amino acid levels in mouse liver after four weeks of vehicle or IMT treatment. Chow Vehicle, Chow IMT and HFD Vehicle n = 8 mice per group, HFD IMT n = 6 mice per group. All data are presented as mean ± SEM. Statistical significance was assessed by a two-way ANOVA with Tukey’s test for multiple comparisons. P-values are shown in the figure.

